# Understanding the physical determinants of pressure denaturation: Exploring the unfolding pathways of Yfh1 at different temperatures with High-Pressure NMR

**DOI:** 10.1101/2024.07.15.603545

**Authors:** Christian Roumestand, Erika Dudas, Rita Puglisi, Antonino Calió, Philippe Barthe, Piero Andrea Temussi, Annalisa Pastore

## Abstract

Proteins unfold under different environmental insults, among which are heat, cold, high pressure and chaotropic agents. Understanding the mechanisms that determine unfolding under each of these conditions is an important problem that directly relates to the physical forces that determine the three-dimensional structure of a protein. Here, we studied a residue-specific description of the unfolding transitions of the marginally stable yeast protein Yfh1 using high-pressure nuclear magnetic resonance. We compared the cold, heat and pressure unfolded states and demonstrated what has up to now been only a hypothesis: the pressure-unfolded spectrum shares features in common with that at low but not at high temperature and room pressure, suggesting a tighter similarity of the mechanisms and a similar role of hydration in these two processes. By exploring the phase diagram of the protein and mapping unfolding onto the three-dimensional structure of the protein, we also show that the pressure-induced unfolding pathways at low and high temperatures differ, suggesting a synergic mechanism between pressure- and temperature-induced denaturation. Our observations help us to reconstruct the structural events determining unfolding and distinguish the mechanisms that rule the different processes of unfolding.

## Introduction

The study of protein unfolding, which corresponds to the process of loss of tertiary structure acquired after the amino acid chain has come out from the ribosome, has fascinated and keeps fascinating generations of scientists because its elucidation holds the promise of understanding the forces that determine protein structure formation. However, structure may be lost, both *in vivo* and *in vitro*, in several different ways, so that speaking about a unique unfolded structure is utterly meaningless (Pastore and Temussi, 2022). In nature, unfolding can be triggered by loss/gain of post-translational modifications, mutations, or any imbalance in the environmental conditions (*e*.*g*. changes in pH, protein concentration, crowding, confinement, etc.). *In vitro*, unfolding can be triggered by changes in temperature, pH, solvent composition, and/or pressure. Among these causes, pressure-induced denaturation is probably the process relatively less studied, possibly because of the non-trivial experimental problems posed by pressurization. The process remains nevertheless extremely interesting both because some proteins do experience unusually high pressure in nature (*e*.*g*. proteins from organisms that live in the deep seas) and because the transition may involve atomic forces probably less dominant in other unfolding processes but not for this less important. Additionally, high pressure presents the advantage to move cold denaturation, an important unfolding transition in principle experienced by all proteins but most often hindered in practice by water freezing (Privalov, 1990), to observable temperatures (Smeller, 2002). Together, these considerations call for a more attentive and detailed study of the mechanisms involved in pressure-induced unfolding.

Despite the pioneering work of several researchers (Zhang et al., 1995; Fuentes and Wand, 1998; Peterson and Wand, 2005; Kitahara et al., 2006; Fu et al., 2012; Vajpai et al., 2013; Nucci et al., 2014; Roche and Royer, 2018; Caro et al., 2022), several aspects of pressure-induced unfolding remain unclear. Most studies of protein unfolding caused by pressure show, for instance, that it is necessary to reach high values of pressure (of the order of thousands of bars) before unfolding can be observed in common piezo-tolerant or pressure mesophile organisms. This might be considered at variance with the devastating effects that comparatively less drastic temperature increases may induce in proteins from common thermally mesophile organisms. The mechanism of pressure-unfolding remains unclear. It is also important to understand what happens during the long lag phases (between 1 and up to several kbar) that are observed in many of the pressure-induced unfolding processes.

We have recently studied the pressure unfolding of a marginally stable protein, Yfh1 from *S. cerevisiae*, that unfolds at pressures so low that its unfolding does not present the typical lag phase usually observed before a protein loses its fold (Puglisi et al., 2022). Yfh1 is a small protein (123 residues in its mature, full-length form) under several aspects remarkable: when depleted of salt, it is possible to observe its cold denaturation at temperatures above water freezing in solutions under quasi-physiological conditions (278 K or 5°C) (Pastore et al., 2007). The high temperature unfolding is also relatively low (around 308 K or 35°C). Also at room temperature, Yfh1 is in an equilibrium between folded and unfolded forms, with a population of the unfolded form around 30%. Addition of even small quantities of salts causes disappearance of the unfolded form (Vilanova et al., 2014). The Yfh1 fold consists of two N- and C-terminal helices that pack on the same side against a 5-7 strand β-sheet depending on the ortholog (Musco et al., 2000). Two interesting features were demonstrated to play an important role in Yfh1 stability. First, the C-terminus of Yfh1 is shorter as compared to other orthologues (*e*.*g*. it is three residues shorter than the bacterial orthologue and twelve residues shorter as compared to the human protein) (Adinolfi et al., 2004). When present, this secondary structure element inserts in between the two terminal helices protecting the hydrophobic core. Accordingly, the melting heat-induced temperatures at low salt of the yeast, bacterial and human proteins go from 308 K, to 328 K, up to 338 K (35°C, 55°C, 65°C) respectively, correlating with the C-terminus length. In Yfh1, where the extension is absent, the core is more easily accessible. In fact, we have demonstrated that shortening the human protein causes a major destabilization, whereas lengthening the yeast protein leads to stabilization (Adinolfi et al., 2004). A second feature is that Yfh1, but not other orthologues, contains four negative residues on the first and second strands (Sanfelice et al., 2014). This quadrilateral of negative charges causes significant electrostatic frustration which leads to cold denaturation under conditions in which hydrophobic forces are weaker: mutation of only one of these residues to a neutral hydrophilic group leads to the shift of cold denaturation at temperatures below the water freezing point while not altering significantly the high-temperature melting point (Sanfelice et al., 2015). Having a system of which we understand in detail the elements that determine protein stability makes Yfh1 a precious tool which we have extensively exploited to probe the mechanisms that determine protein unfolding.

In a previous study, we determined the phase diagram of Yfh1 unfolding as a function of pressure (1-5000 bar) and temperature 278-313 K (5-40°C), both in the absence and in the presence of fold stabilizers using the intrinsic fluorescence of the two tryptophan residues of Yfh1 (Puglisi et al., 2022). We demonstrated that Yfh1 has a remarkable sensitivity to pressure: 50% unfolding occurs already at pressures around 100 bar at room temperature. For comparison, more thermodynamically stable globular proteins, such as the immunoglobulin-like module of titin I27 or a hyper-stable variant of Staphylococcus nuclease, need pressures as high as 2-3 kbar to unfold in mild concentrations of guanidinium chloride, showing an incredible resilience (Herrada et al. 2018; Roche et al., 2012).

In the present study, we exploited these unusual properties to study the pressure-unfolding pathways of Yfh1 as a function of temperature by nuclear magnetic resonance (NMR). This technique is almost unique in providing residue-specific information on the folding/unfolding pathway of a given protein (Pastore and Temussi, 2023; Roche et al., 2017). We show that the pressure-unfolded NMR spectrum shares closer features with the low but not the high temperature-unfolded spectrum, suggesting a similar mechanism. We also explored the phase diagram of the protein, and, thanks to the residue-specific information provided by high-pressure NMR, we could reconstruct the pathway that determine unfolding. This evidence provides us with a detailed model of how unfolding occurs under different conditions. We can thus conclude that, as compared to fluorescence, NMR provides complementary information and allows a more thorough understanding of the processes occurring during protein unfolding.

## Materials and Methods

### Sample Preparation

The recombinant Yfh1 protein was produced as previously described (Adinolfi et al, 2002). In short, the ^15^N-labeled protein was expressed in *Escherichia coli* BL21-(DE3) cells grown at 314 K (37 °C) in minimal medium using ammonium sulphate as the sole source of nitrogen, and induced by addition of 0.5 mM IPTG for 2 h. After cell harvesting by centrifugation, the cells were re-suspended in Tris-HCl buffer containing a complete EDTA protease inhibitor cocktail tablet (Roche) and lysed by sonication. The soluble protein was purified by two ammonium sulphate precipitation steps at 40% cut to precipitate contaminating proteins and a 65% cut to precipitate Yfh1. The protein was dialysed and further purified by anion exchange chromatography using a Pharmacia Q-Sepharose column with a gradient from 0 to 1 M NaCl, followed by a Pharmacia phenyl-Sepharose column with a decreasing 1 M ammonium sulphate gradient. The final samples were dialysed in 20 mM HEPES at pH 7.0. No additional salt was added since we had previously demonstrated that even small quantities of salt appreciably stabilize the protein (Adinolfi et al., 2004; Vilanova et al., 2014).

### Protein unfolding monitored by high-pressure NMR Spectroscopy

A protein sample with about 0.5 mM concentration of ^15^N-labeled Yfh1 in 20 mM HEPES at pH 7.0, with 2 mM DTT and containing 5% (v/v) D_2_O for the lock, was used on a 5/3 mm O.D./I.D. ceramic tube (330 µl of sample volume) from Daedalus Innovations (Aston, PA, USA). Hydrostatic pressure was applied to the sample directly within the magnet using the Xtreme Syringe Pump also from Daedalus Innovations. One ^1^H and two-dimensional [^1^H,^15^N] HSQC spectra were recorded on a Bruker AVANCE III 600 MHz spectrometer (standard ^1^H-^15^N double-resonance BBI probe), in the range 278-303 K, although in the final analysis we used only those at 283-303 K. At each temperature, the pressure was varied from 1 bar to 1000 bar by steps of 50 bar. Each pressure step lasted 2 h, starting with 1 h relaxation time, to allow the folding/unfolding reaction to reach full equilibrium, followed by a 1D spectrum (32 scans with 10 s of relaxation time, to ensure the complete relaxation of the methyl protons between each scan) and a 2D [^1^H,^15^N] HSQC (8 scans for each of the 128 complex points in the indirect dimension). The relaxation time preceding the recording of the experiments was estimated from a series of 1D NMR experiments recorded after a 300 bar P-Jump, following the exponential growth of the resonance band corresponding to the methyl groups in the unfolded state of the protein (Dubois et al, 2020). Reversibility of unfolding was checked by comparing 1D and 2D [^1^H,^15^N] HSQC spectra recorded at the end of the series of experiments, after returning at 1 bar, with the spectra recorded at 1 bar before pressurization.

^1^H chemical shifts were directly referenced to the methyl resonance of DSS (2,2-Dimethyl-2-silapentane-5-sulfonate, sodium salt), while ^15^N chemical shifts were referenced indirectly to the absolute frequency ratios ^15^N/^1^H = 0.101329118. The water signal was suppressed by the WATERGATE pulse sequence (Piotto et al., 1992). The assignment of the amide resonances of Yfh1 was retrieved from the BioMagResBank (http://www.bmrb.wisc.edu, entry number 19991). Data processing and analysis of the HSQC experiments were performed using GIFA (Pons et al., 1996). At each temperature, the cross-peak intensities for the folded species were measured at each pressure and then fitted with a sigmoidal curve characteristic of a two-state equilibrium:

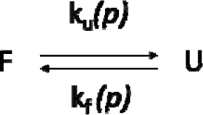

Where F and U correspond to a given residue (identified through its amide cross-peak) sitting in a folded or unfolded state respectively, in equilibrium with an equilibrium constant K_eq_ of:

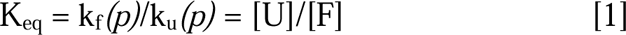

where k_f_*(p)* and k_u_*(p)* stands for the folding and unfolding rate constants at a given pressure *p*. K_eq_ can be also expressed from the Boltzmann equation as:

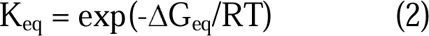

where the free energy change ΔG can be expressed as a Taylor expansion, truncated at the second order term:

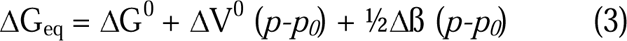

ΔG_eq_ and ΔG^0^ are the Gibbs-free energy changes from F to U at pressure *p* and *p*_*0*_ (*p*_*0*_ = 1 bar), respectively; ΔV^0^ is the variation of partial molar volume; Δβ is the variation in the compressibility coefficient, R is the gas constant, and T is the absolute temperature. It has been shown that, for proteins, the difference in compressibility between native and denatured states is negligible (Ravindra, 2003). Thus, the expression of ΔG_eq_ simplifies to:

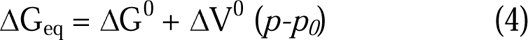

Using amide cross-peak intensities I as the observables, the equilibrium constant can be written as:

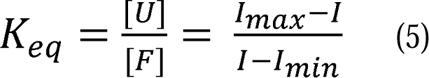

where, for each temperature, I is the cross-peak intensity of the folded species measured at a given variable pressure, and I_max_ and I_min_ correspond respectively to the intensities of the same cross-peak in the fully native (low pressure) and in the unfolded states (high pressure). Combining this equation with equations (2) and (4) gives the characteristic equation for a two-state equilibrium:

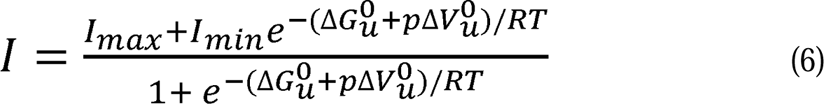

In the fitting, I_max_, I_min_, 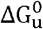 and 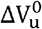 values were left as floating parameters. Once the 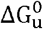 and 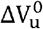 values were obtained, the residue-specific curves were normalized (I_max_= 1 and I_min_ = 0 in Eq. 6) to give the native fraction of the protein as a function of pressure.

### Calculation of the ΔH_m_, ΔC_p_, and T_m_ values at ambient pressure

Assuming that the difference in heat capacity, Δ*C*_*p*_, between native and unfolded state is temperature independent, ΔG^0^ at constant pressure depends on temperature as described by the modified Gibbs-Helmholtz equation:

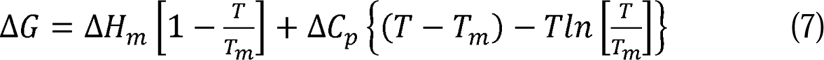

where *T*_*m*_ is the temperature at the midpoint of the unfolding transition, and Δ*H*_*m*_ is the unfolding enthalpy change at *T*_*m*_. The curve corresponding to this equation is known as the stability curve of the protein. The thermodynamic parameters Δ*H*_*m*_, Δ*C*_*p*_, and *T*_*m*_ were obtained by fitting to Eq. 7 the average values 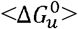 obtained by averaging the residue-specific values of 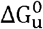 measured at each temperature for each residue, through their corresponding residue-specific pressure denaturation curve. Nonlinear fitting of equations by experimental data was carried out using the Levenberg-Marquardt algorithm.

### Calculation of the contact maps

According to a method previously developed (Roche et al., 2012), we defined the probability of contact for each pair of residues, *P*_*i,j*_, as the geometric mean of the fractional probability of the two residues at a given pressure using the relation (Fossat et al., 2016):

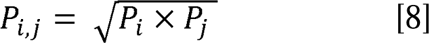

where the fractional probability *P*_*i*_ or *P*_*j*_ correspond to the probability to find residue *i* or *j* in the native state at a given pressure for a given temperature. These fractional probabilities are obtained directly from the normalized residue-specific denaturation curves obtained for each residue (Dubois et al., 2020). Using CMview (Vehlow et et., 2011; http://www.bioinformatics.org/cmview/) with a generous cut-off distance threshold of 9.5 Å, a total of 662 (non-sequential) contacts between the Cα of the 123 residues of the protein could be measured from the AlphaFold model of Yfh1 (Jumper et al., 2021). Of them, 250 contacts concern the 67 residues for which residue-specific denaturation curves could be obtained. We then plotted the evolution of the number of lost contacts as a function of pressure and temperature, assuming that a contact is “lost” when *P*_*i,j*_ < 0.5.

## Results

### Modelling a reference 3D structure of Yfh1

Although the structures of several Yfh1 orthologues from different species have now been solved, the structure of the yeast protein remains determined only at low resolution. Of the seven entries available in PDB, 2GA5 is an NMR structure which however has serious distortions of the geometry caused by assignment misinterpretation: as previously demonstrated (Vilanova et al., 2014), the authors mistraced sequential assignment, mixing resonances from the folded and the unfolded states. This caused strain and a poor geometry. The other six structures, solved by crystallography at *ca*. 3 Å resolution, are those of a mutant in which a well conserved tyrosine (Tyr73) was substituted by an alanine (Söderberg et al., 2011). This mutation enhances the tendency of the protein to form large aggregates in the presence of high concentrations of iron but affects the structure of the N-terminal helix. The NMR and crystallographic structures superpose with a 2.5 Å rmsd (on residues 75 to 172 using the numbering of the mature protein and of the 3OEQ entry), with maximal differences in the orientation of the C-terminal helix and in the structure of the 6^th^ and 7^th^ putative ß-strands. To clarify the issue, we used AlphaFold to generate a model that could be consistent with the structures of all Yfh1 orthologues. The AlphaFold vs 2.2 software used for modelling yielded a model with a high degree of confidence: only eight out of the 123 residues of Yfh1 had a pLDDT (predicted local distance difference test) score lower than 0.9, all located at the N-terminal extremity of the molecule, which has been demonstrated to be intrinsically disordered and flexible (Popovic et al., 2015) (**Figure 1**). This model superposes with the crystal structure with a 1.1 Å rmsd in the globular region (residues 75-172), as compared to a 2.2 Å rmsd with the NMR structure. However, although a better superposition is observed for the ß-sheet between the X-ray structure (3OEQ) and the AlphaFold model, the lengths of the ß-strands are not always the same. The X-ray structure also lacks, due to the structural distortions induced by the Tyr73Ala mutation, the first two helical turns of the N-terminal helix (L70-E76) that are present both in the AlphaFold model and in the NMR structure (2GA5). The crystal structure lacks the two last residues (S172, Q173), yielding to a shortening of the C-terminal helix where an additional turn is observed both in the NMR structure and on the AlphaFold model. For these reasons, we decided to consider the AlphaFold model as the reference for any further analysis.

**Figure 1.**
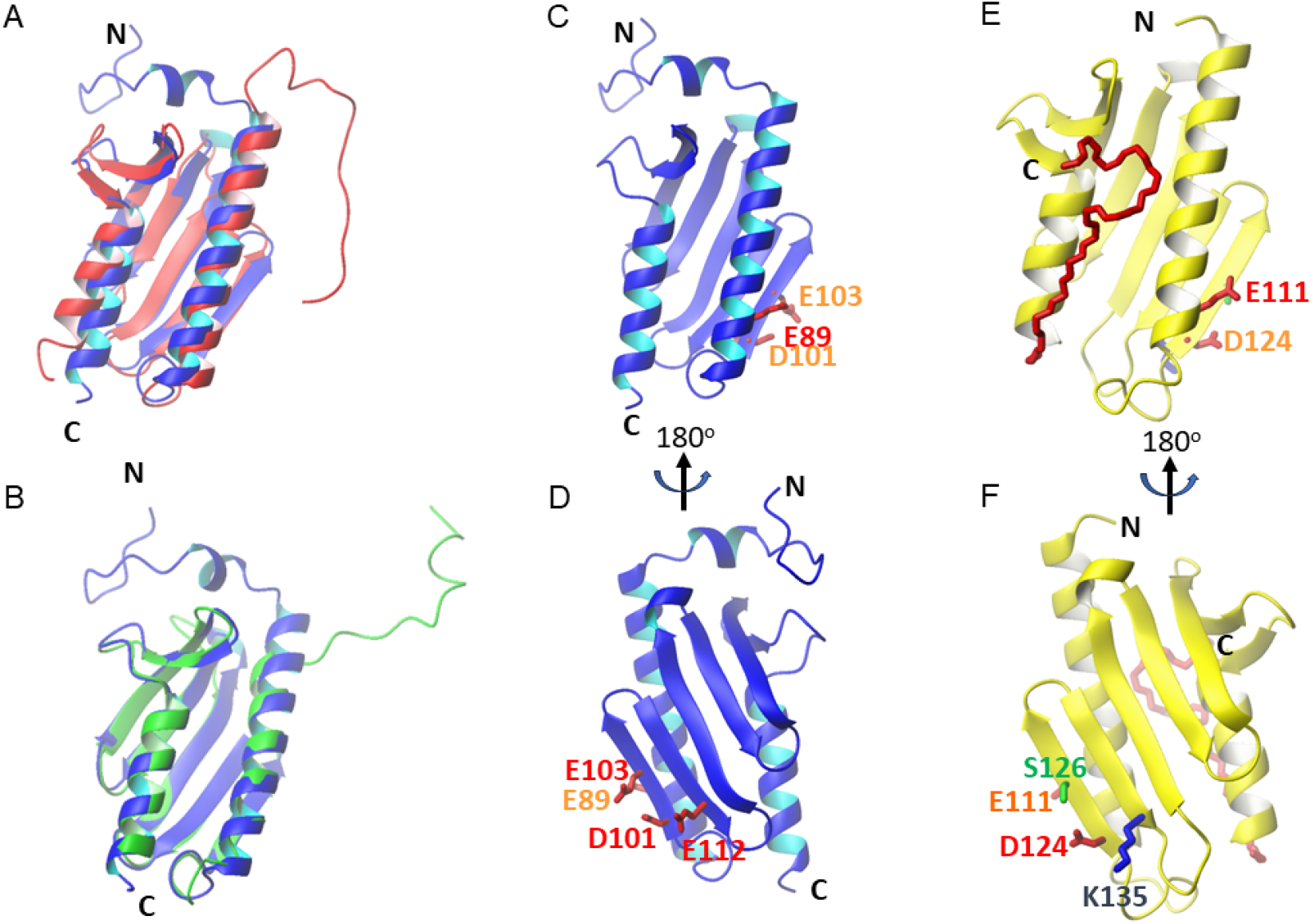
Comparison between the experimental structures and the AlphaFold model of Yfh1. Superimposition of the 3D structures (ribbon representations) of the AlphaFold model of Yfh1 (blue ribbon) with **A**) the corresponding NMR solution structure (2ga5) and **B**) the X-ray structure (3eoq) (green ribbons). C) and D) Ribbon representation of the AlphaFold model of YFH1 in two different orientations differing by a 180° rotation around the y axis, highlighting the negatively charged cluster that promotes electrostatic frustration and the C-short terminus. E) and F) For comparison the structure of human frataxin (1egk) on which the residues corresponding to the electrostatic cluster are indicated and the much longer C-terminus that inserts in between the two helices and protects the hydrophobic core. The side chains of negatively charged residues are indicated in red, positive residues in blue and non-charged residues in green.

### Yfh1 undergoes pressure denaturation under modest pressures

To explore the phase diagram of Yfh1 and get residue-specific information of the unfolding pathways of Yfh1, we recorded 1D ^1^H and 2D [^1^H,^15^N] HSQC spectra in the pressure range of 1-1000 bar and between 278 and 303 K with a 5 K interval to analyse Yfh1 pressure unfolding at the residue level (Roche et al., 2017; Roche et al., 2019; Dubois et al., 2020). This range of pressure was chosen because previous high-pressure fluorescence spectroscopy studies of Yfh1 had demonstrated that the protein has the midpoint of unfolding between 100-200 bar (Puglisi et al., 2022). Note that the spectra at 278 K were recorded but not further analysed: at this temperature, the protein is already largely unfolded at atmospheric pressure and the remaining resonances corresponding to the native fraction are severely broadened. For reference, we also compared the behaviour of the 1D peak areas versus temperature at atmospheric pressure with previous similar plots (Pastore et al., 2007) to assess reproducibility, and observed an excellent qualitative agreement (**Figure S1**).

The 2D [^1^H,^15^N] HSQC spectra at 1 bar, recorded at different temperatures and low ionic strength, were of excellent quality, with well-dispersed resonances and several high- and low-frequency resonances which demonstrated that the protein is mainly folded in the range of temperatures 283-303 K (**Figure 2**). As usually observed, the spectra at increasing pressures showed progressive attenuation of the resonances from the folded species. Conversely, the resonances of the unfolded species became increasingly intense and peaks originally less intense or masked by other more intense resonances appeared. The presence of the equilibrium between folded and unfolded species in a slow exchange regime in the NMR timescale is evident from the co-existence in the HSQC spectra of well dispersed resonances in co-presence with a number of other peaks that change their intensity at different temperatures that are diagnostic of the unfolded species. An example is the cross-peak at 10.2 ppm/129 ppm (^1^H/^15^N) that corresponds to the indole resonances of the two Trp residues in the unfolded species. These resonances, that are present also in the spectrum at 1 bar and at the temperature of maximal stability, disappear upon addition of also minute concentrations of salt (Vilanova et al., 2014).

**Figure 2.**
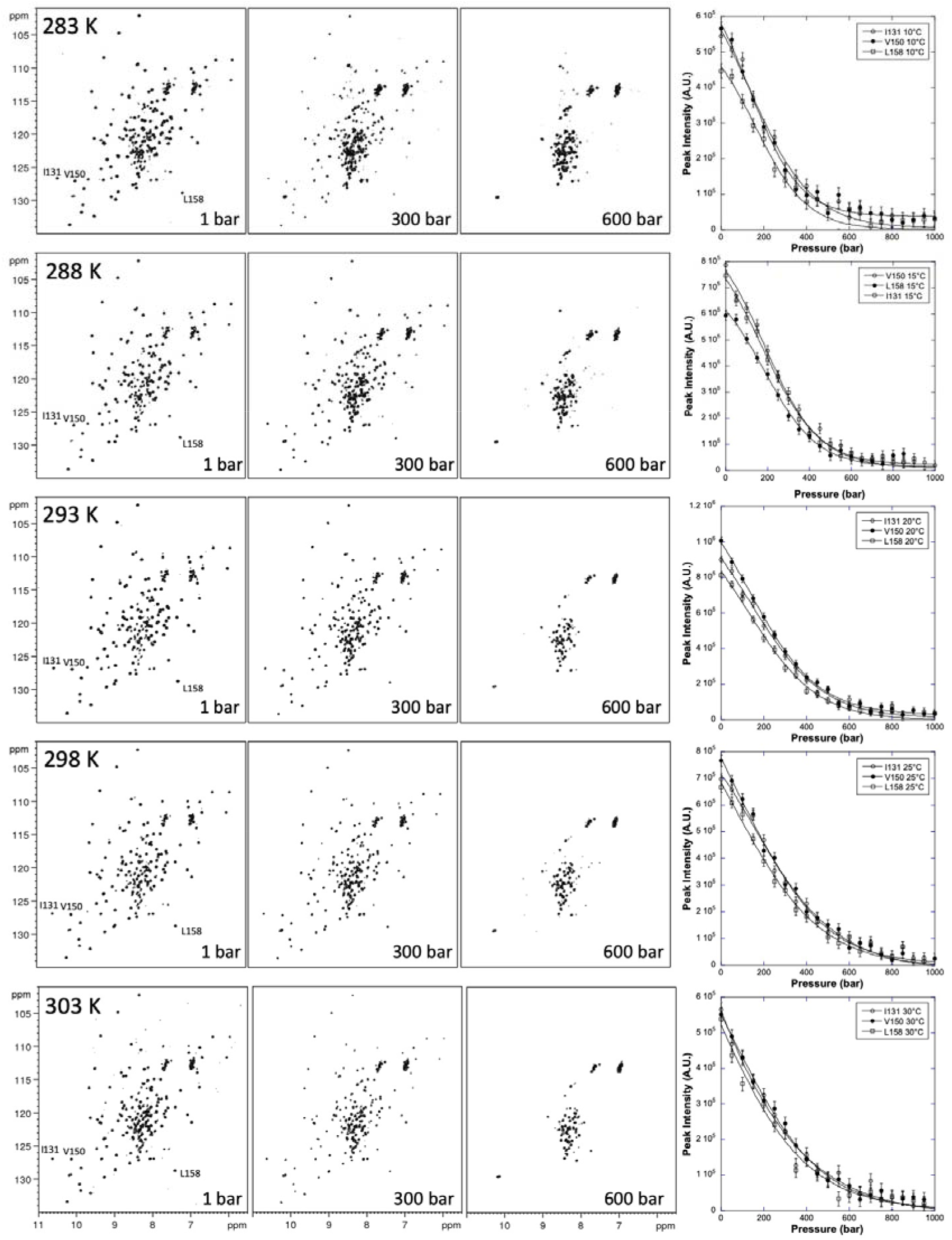
NMR monitored high-pressure unfolding of Yfh1. [^1^H,^15^N] HSQC spectra recorded at 283 K, 288 K, 293 K, 298 K and 303 K (from top to bottom, as indicated). At each temperature, spectra at 1, 300 and 600 bar are displayed from left to right. The rightmost panels report the overlays of residue-specific denaturation curves for three representative residues (I131, V150, and L1584, labelled on the corresponding HSQC at 1 bar) obtained from the fits of the pressure-dependent sigmoidal decrease of the corresponding residue cross-peak intensities in the HSQC spectra with Eq. 6.

A total of 67 resonances (corresponding to 59% of the 114 non-proline residues) had no overlap with other peaks at any of the five temperatures and displayed cross-peaks of sufficient intensity at atmospheric pressure to be accurately fit by a two-state model, providing an appreciable number of local probes for the description of Yfh1 unfolding.

Above ca. 400 bar, we observed nearly complete disappearance of the well-dispersed species, suggesting almost complete unfolding of the protein. Some residual structure remained visible also at 600 bar at the noise level, but the relative ratio accounts for less than 2% of the initial intensity. The high-pressure spectra of the pressure-unfolded species exhibited a similar collapse of the resonances at all recorded temperatures, indicating that the primary denaturing agent is pressure (**Figure 2**).

After reaching 1000 bar, the sample was returned to 1 bar in two steps (500 and 1 bar). The spectra collected before and after pressurization were superimposable within 10% differences, demonstrating the almost complete reversibility of the process. This observation is fully in line with our previous experience with Yfh1, that is a protein altogether with relatively low tendency to aggregate (covalently and non-covalently).

### The cold and pressure unfolding states share some closer features

In previous studies, we had made a detailed comparison between the chemical shifts of the amide protons of the unfolded states at low and high temperatures (Adrover et al., 2010 and 2012). We had found that the amide secondary chemical shifts, that is the difference between the experimental values from a given residue and the random-coil values recorded at 298 K for model tetrapeptides (Wishart et al., 1995), had consistent different and opposite signs: the values measured at low temperature were negative, implying de-shielding, whereas those at high temperature were positive, indicating shielding effects (Adrover et al., 2010). We had explained these results by a different degree of hydrogen bonding of the amides with water, reflecting a higher degree of hydration at low temperature.

Here, we compared the spectra of the unfolded states at 15°C, where the protein reaches its maximum of stability, and at high pressure (800 bar) with those obtained at high and low temperatures and atmospheric pressure (Adrover et al., 2010) (**Figure 3**). Several observations could be extracted from the comparison: first, a similar level of collapse of the spectrum is evident in all three conditions, indicating population of mostly unfolded states. Second, the spectrum at high pressure looks overall closer to that at low temperature and 1 bar, while the high temperature spectrum appears more broadened, likely indicating a higher degree of conformational exchange between species. Third, the spectrum at high pressure exhibits an overall low-field shift which suggests a de-shielding similar to that at low temperature and 1 bar. This can be seen by comparing the overall position of the spectra as compared to an ideal oval plotted in the same position. The shift is also evident for the well-isolated resonances of the only two tryptophan residues in the sequence, one buried and one exposed that are visible around 10.2 ppm and 129 ppm in the proton and nitrogen dimensions respectively in all denatured spectra (**Figure 3**). Due to sequential effects, the two peaks are not completely superimposable indicating residual local level of asymmetric environments, but they are clearly downfield shifted in the cold denatured and high-pressure spectra by ca. 0.2 ppm as compared to high temperature.

**Figure 3.**
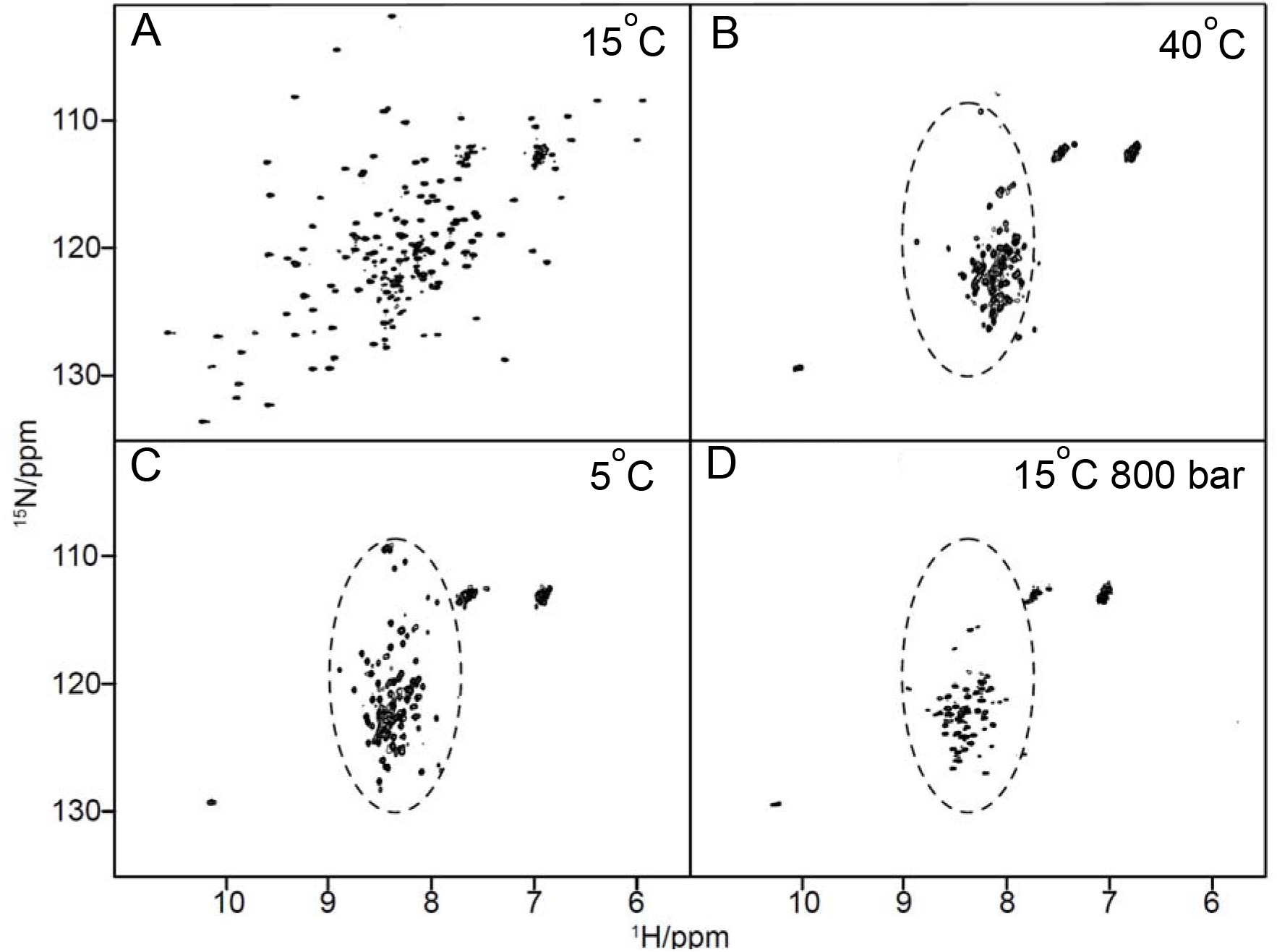
Comparison of the HSQC spectra of Yfh1 under different conditions. A) Spectrum recorded at the temperature of maximal stability of the protein and room pressure. The spectrum has excellent dispersion and all the features typical of a folded protein. B) Unfolded spectrum at 313 K and room pressure. C) Unfolded spectrum at 278 K and room pressure. D) Unfolded spectrum at 288 K and 800 bar. A dotted oval is drawn in the same region of the various unfolded spectra which roughly corresponds to the area span by all peaks in the cold denatured spectrum. The three spectra show a similar collapse of the resonance dispersion, but the heat denatured is noticeably up-shifted as compared to the cold denatured spectrum as previously analysed (Adrover et al., 2010 and 2012). This indicates a different level of hydration that is higher at low temperature. The pressure-denatured spectrum has overall features closer to the cold denatured one.

Altogether, these observations can be explained by a similar degree of hydrogen bonding strength of the amides with water at low temperature and high pressure, indicating a higher similarity between these two unfolded states as compared to the high temperature one.

### Exploring the unfolding pathways of Yfh1 at different temperatures by High-Pressure NMR

Fitting the cross-peak intensities with the corresponding two-state equation (Eq. 6) provided residue-specific values for the apparent free energy 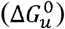 of unfolding (**Figure 4**), which reports on the protein stability, and the apparent volume change 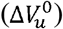 of unfolding (**Figure 5**), which corresponds to the volume difference between the folded and unfolded states of the protein.

**Figure 4.**
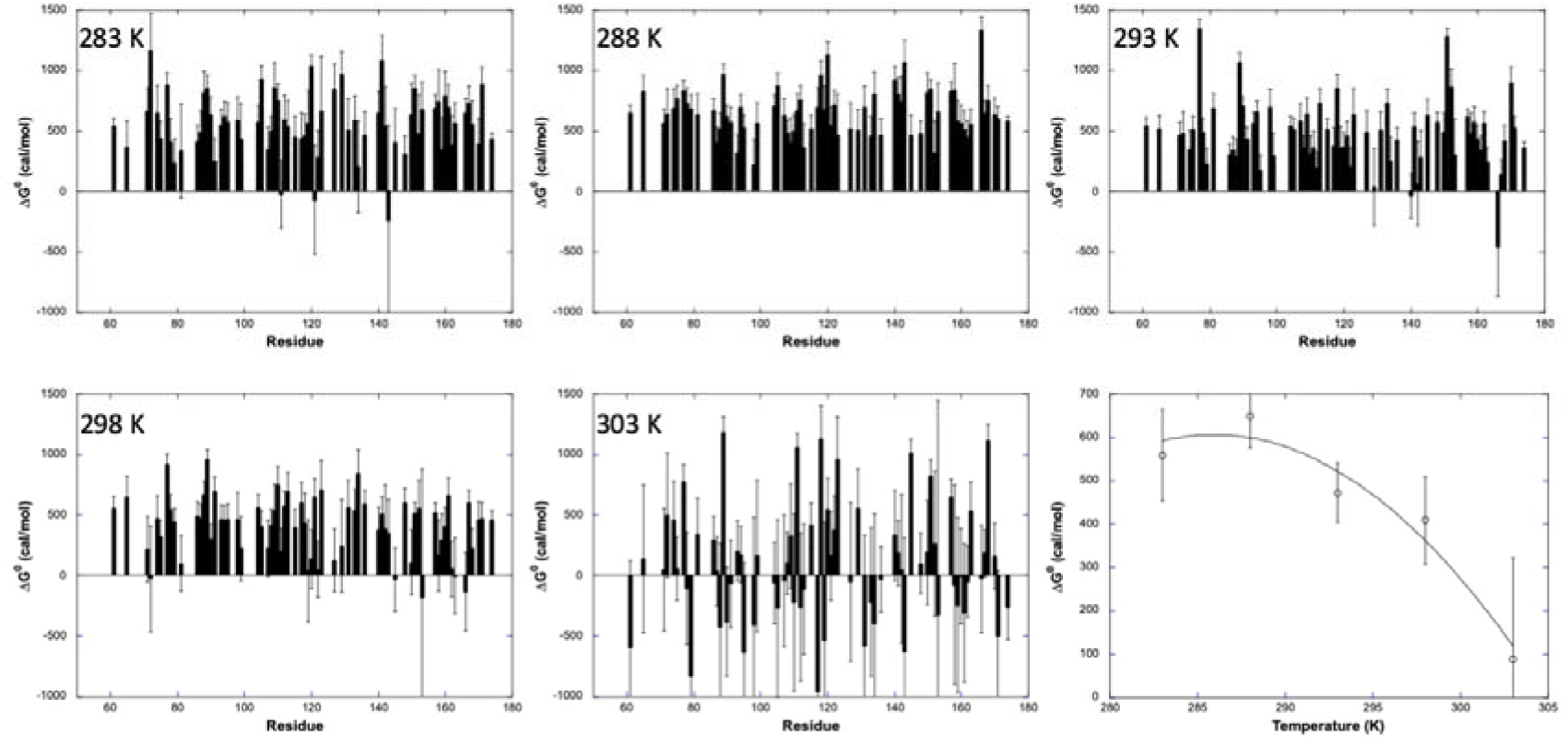
Apparent Residue-Specific difference of Gibbs Free Energy of Unfolding 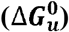. values measured for Yfh1 at 283 K, 288 K, 293 K, 298 K and 303 K (as indicated) from residue-specific pressure denaturation curves. The residue numbering used in the figure corresponds to the one deposited for the X-ray structure (3EOQ). The last panel displays the averaged values of 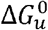 at each temperature versus the temperature, fitted with the Gibbs-Helmotz equation (Eq. 7).

**Figure 5.**
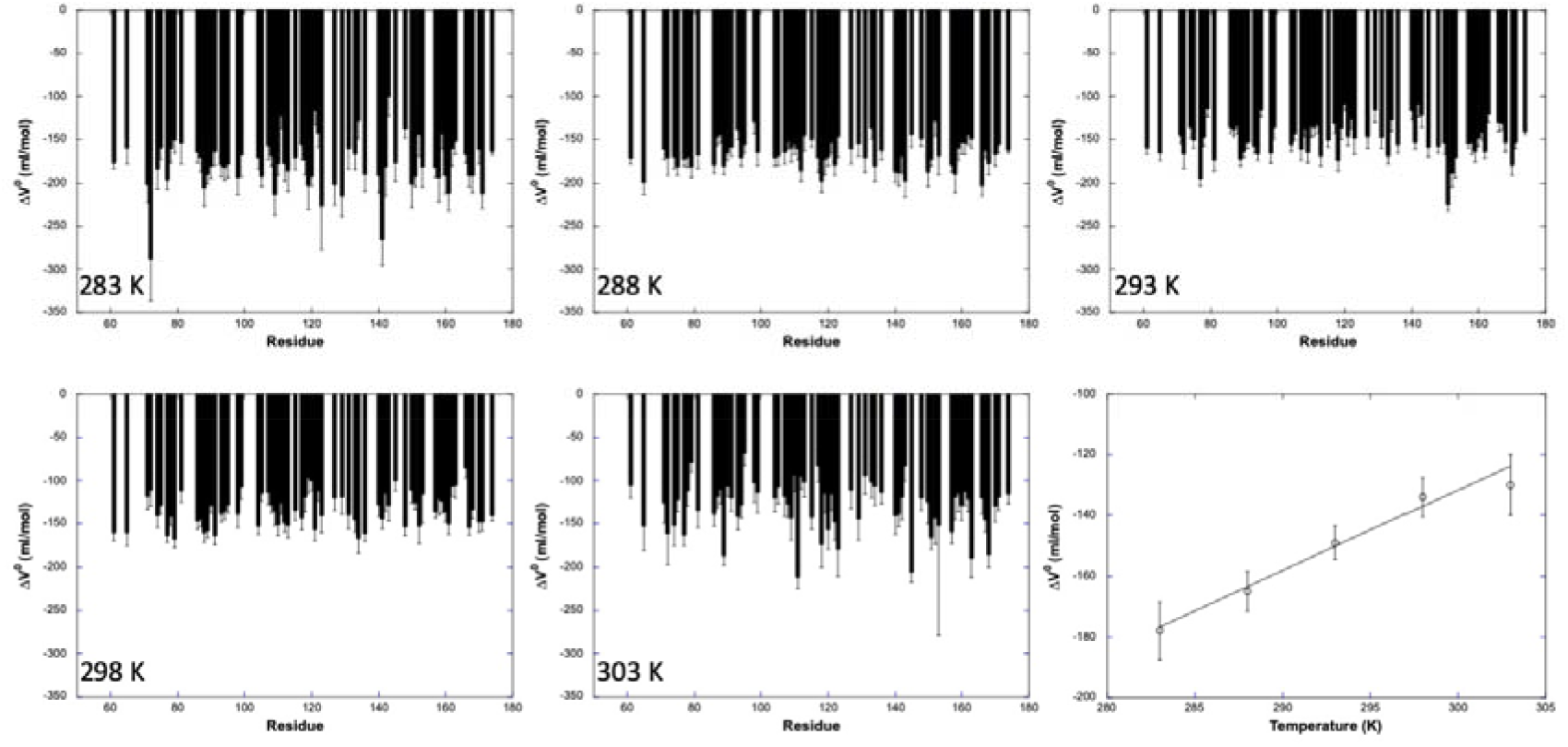
Apparent Residue-Specific Volume Change of Unfolding 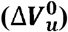. values measured for Yfh1 at 283 K, 288 K, 293 K, 298 K and 303 K (as indicated) from residue-specific pressure denaturation curves. The residue numbering used in the figure corresponds to the one deposited for the X-ray structure (3EOQ). The last panel displays the linear fit of the averaged values of 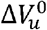 at each temperature versus the temperature.

The two-state model was adequate to fit most of residue-specific unfolding curves, even though the absence of points describing the upper plateau (I_max_) of the sigmoid, due to partial unfolding already at 1 bar, yielded significant error bars for the apparent residue-specific free energy values 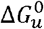 and, to a lesser extent, for the residue-specific 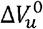 values (**Figure 4 and 5**). This is especially true for experiments recorded at 303 K, temperature at which several residues exhibit negative values of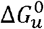, meaning that these residues are in an unfolded conformation at atmospheric pressure for more than half of the protein population.

Yfh1 displays low stability, that appears to be maximal around 288 K, with an average value for the apparent free energy of unfolding 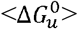 of 649 ± 145 cal/mol, and significantly decreases at higher and lower temperatures (**Table 1**). These values are remarkably low when compared to what usually observed for other small globular proteins (Smeller, 2002), that often needed the presence of guanidinium chloride to observe pressure unfolding within a feasible pressure range (Roche et al., 2012; Herrada et al., 2018; Saotome et al., 2019, Lahfa et al., 2023). The dependence of 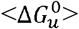 on temperature exhibited a concave profile with a maximum around 288 K (**Figure 4**). Fitting the temperature dependence of 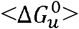 at atmospheric pressure to the Gibbs-Helmotz equation (Eq. 7) yielded the averaged thermodynamic parameters ΔC_p_ of unfolding (1 ± 0.7 kcal/mol.K), T_m_ (305 ± 4 K) and ΔH_m_, (19 ± 7 kcal/mol) (**Figure 4 and Table 2**) which are in excellent agreement with previous thermal denaturation studies (Pastore et al., 2007; Martin et al., 2008).

**Table 1.** Average residue-specific apparent free energy 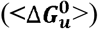 and volume 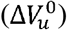 of unfolding values measured at equilibrium and at atmospheric pressure and different temperatures. The values are compared to the equivalent ones obtained from fluorescence spectroscopy (Puglisi et al., 2022).

| Temp<br>(°C/K) | $\langle \Delta G_u^0 \rangle$ (kcal/mol) | $\Delta V_u^0$ (ml/mol) | $\langle \Delta G_u^0 \rangle$ (kcal/mol) | $\Delta V_u^0$ (ml/mol) |
| --- | --- | --- | --- | --- |
|  | Present study (NMR) |  | Puglisi et al., 2022 (Fluorescence) |  |
| 5/ 278 | ND | ND | $-0.20 \pm 0.02$ | $-87 \pm 2$ |
| 10/ 283 | $0.559 \pm 0.212$ | $-178 \pm 19$ | $-0.03 \pm 0.02$ | $-90 \pm 4$ |
| 15/ 288 | $0.649 \pm 0.145$ | $-165 \pm 13$ | $0.09 \pm 0.05$ | $-90 \pm 2$ |
| 20/ 293 | $0.472 \pm 0.138$ | $-149 \pm 11$ | $0.22 \pm 0.07$ | $-83 \pm 3$ |
| 25/ 298 | $0.409 \pm 0.202$ | $-134 \pm 13$ | $0.17 \pm 0.02$ | $-81 \pm 2$ |
| 30/ 303 | $0.089 \pm 0.264$ | $-130 \pm 20$ | $-0.08 \pm 0.02$ | $-70 \pm 1$ |
| 40/ 313 | ND | ND | $-0.19 \pm 0.05$ | $-59 \pm 1$ |

**Table 2.** Comparison of the thermodynamic parameters for cold and heat unfolding of Yfh1 at atmospheric pressure obtained in the present study with values previously reported.

| | $\Delta H_m$ (kcal mol<sup>-1</sup>) | $\Delta C_p$ (kcal K<sup>-1</sup> mol<sup>-1</sup>) | $T_c$ (°C) | $T_m$ (°C) |
| --- | --- | --- | --- | --- |
| Present study | $19 \pm 7$ | $1.0 \pm 0.7$ | n.d. | $32 \pm 4$ |
| Pastore et al., 2007 | $21 \pm 2$ | $1.8 \pm 0.1$ | $7.0 \pm 1.5$ | $31 \pm 2$ |
| Martin et al., 2008 | 20.5 | 1.9 | 8 | 29.2 |

A linear decrease with temperature was observed for the average absolute values of the apparent 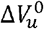 **(Figure 5)**. The slope of this dependence of 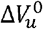 on temperature corresponds to the difference in thermal expansivity, Δα, between the folded and unfolded states of the protein. The estimated value of Δα = 2.7 ± 0.7 ml/mol.K is comparable to those found in the literature (Dubois et al., 2021; Fossat et al., 2016; Dellarole et al., 2015; Rouget et al., 2010). On the other hand, the absolute values of 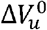 (≈ 150 ml/mol) are substantially greater than those usually found for proteins of comparable size and from those measured on Yfh1 by fluorescence spectroscopy (≈ 90 ml/mol) (Puglisi et al., 2022).

The residue-specific apparent values of 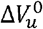 and 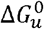 were used to build normalized residue-specific denaturation curves, giving the fraction of folded species for each residue as a function of pressure for each of the five temperatures used in this study (**Figure 6**) (Roche et al.; 2012; Dubois et al., 2020). The average normalized curve was then calculated from the individual residue-specific normalized curves, giving information on the global evolution of the native fraction of the protein during pressure unfolding. Plots of the evolution of the native fraction as a function of the temperature at different pressures (1, 150, 300, 450 and 600 bar) showed a maximum for the native fraction around 288 K (**Figures 6**), consistent with the temperature dependence of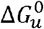. Interestingly, the differences observed between temperatures diminish when increasing the pressure, until merely vanishing at high pressure (600 bar), indicating that the total unfolding of the protein is reached around the same pressure whatever the temperature. This strongly suggests that temperature does not only affect the stability of the protein, but also the cooperativity of unfolding.

**Figure 6.**
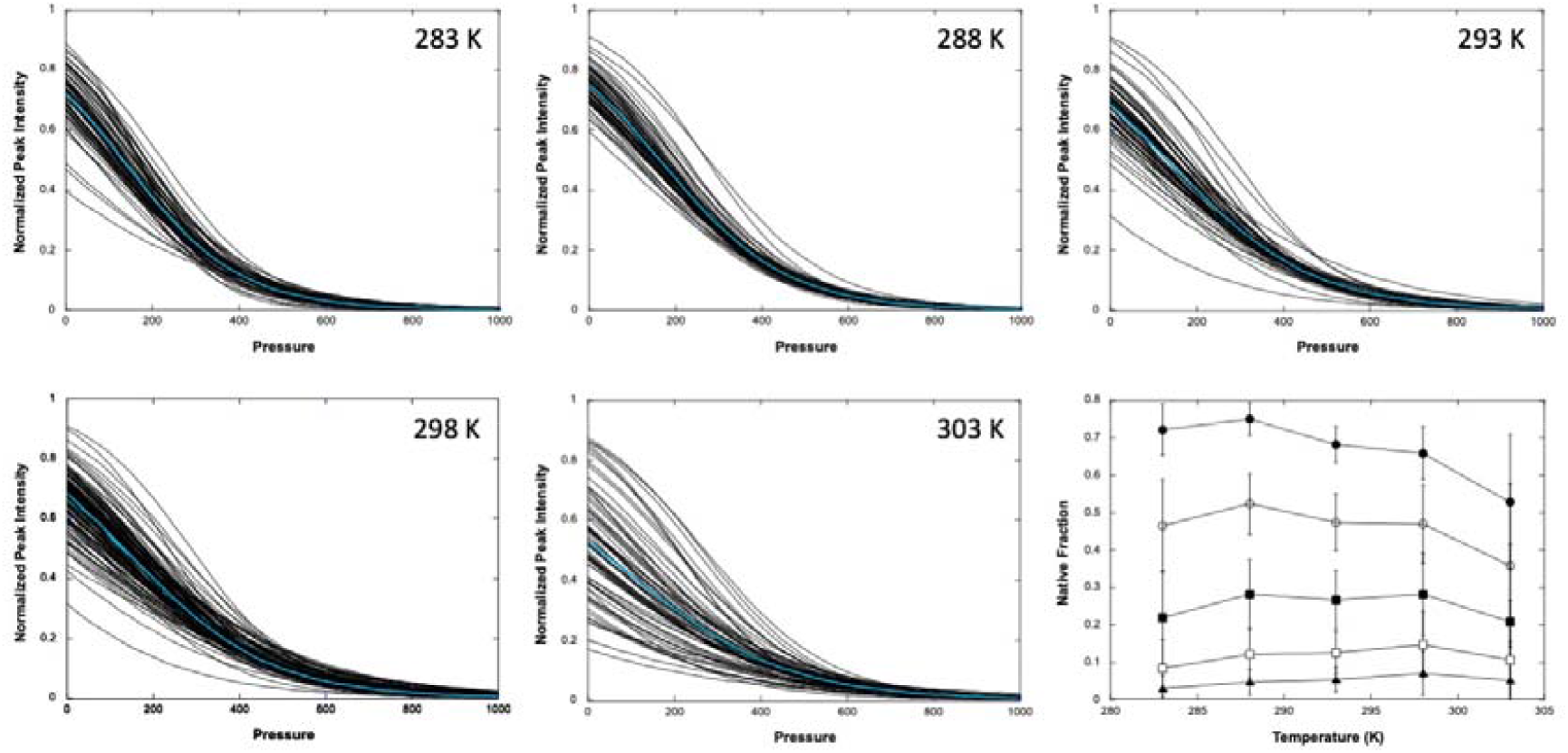
Normalized residue-specific denaturation curves at different temperatures. The bold cyan curves represent the average normalized curve at each temperature. The last panel displays the evolution of the native fraction of the protein (obtained from the average normalized curves) as a function of the temperature at 1 (filled circles), 150 (open circles), 300 (filled squares), 450 (open squares) and 600 bar (filled triangles).

### Evidence of different unfolding pathways at different temperatures

As already reported for other proteins, it is possible to characterize the folding pathway of a protein by mapping on the structure the regions of the protein that progressively become unfolded at increasing pressures following a procedure previously described (Roche et al., 2012). Consistent with the 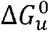 values, the number of lost contacts at 150 bar is much less important at 288 K than at lower or higher temperatures, reflecting the higher stability of the protein at this temperature (**Figure S2**). Virtually all contacts were lost at 200 bar at all temperatures, with partial unfolding of the protein starting at 50 bar at 283 K, 293 K and 298 K. Note that at this pressure, the number of lost contacts is above 60% at 303 K, confirming that the protein is already severely denatured under these conditions. At 288 K, the unfolding transition appears more cooperative, with only 6% of lost contacts at 125 bar, although the protein is almost totally unfolded already at ∼200 bar. Thus, the higher stability of the protein at this temperature is associated with a higher cooperativity of its unfolding transition.

We then built fractional contact maps from the probabilities of contact calculated at the five temperatures (**Figure 7**). At 283 K, few long-range contacts are lost (*P*_*i,j*_ < 0.5) in the ß-sheet at 50 bar, as well as a few long-range contacts in the ß-sheet, as well as a few longrange contacts between the beginning of the ß-sheet and the N-terminal helix. At 100 bar, contact loss concerns more residues in the ß-sheet, and also to long-range contacts between the ß-sheet and the two helices. At this pressure, the N-terminal helix begins to unfold, whereas the C-terminal helix remains mostly unaffected. At 150 bar, almost all contacts are lost. At 288 K, that is close to the maximal stability temperature, unfolding appears very cooperative, starting only at 150 bar, and concerning all the secondary structure elements, as well as the tertiary contacts between them. At 200 bar, virtually all contacts are lost, confirming the higher cooperativity of the unfolding reaction at this temperature. At higher temperatures, contrary to what observed at 283 K, unfolding becomes again less cooperative, and contact loss concerns first the C-terminal helix (50 bar) and then extends to the ß-sheet (100 bar), the N-terminal helix being the last secondary structure element to unfold. At 303 K, almost all contacts are already lost at 50 bar, hampering comparison with the other temperatures.

**Figure 7.**
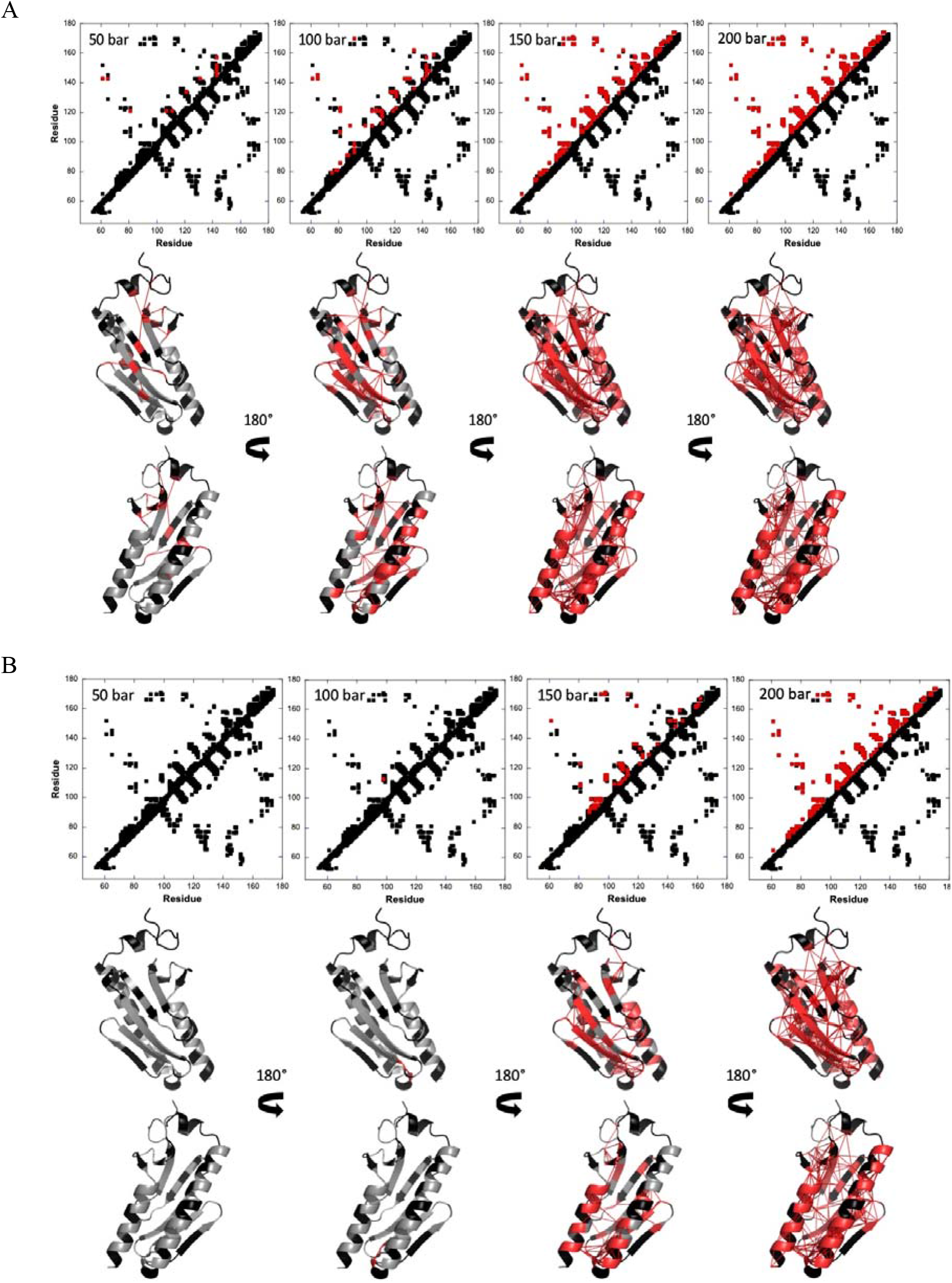
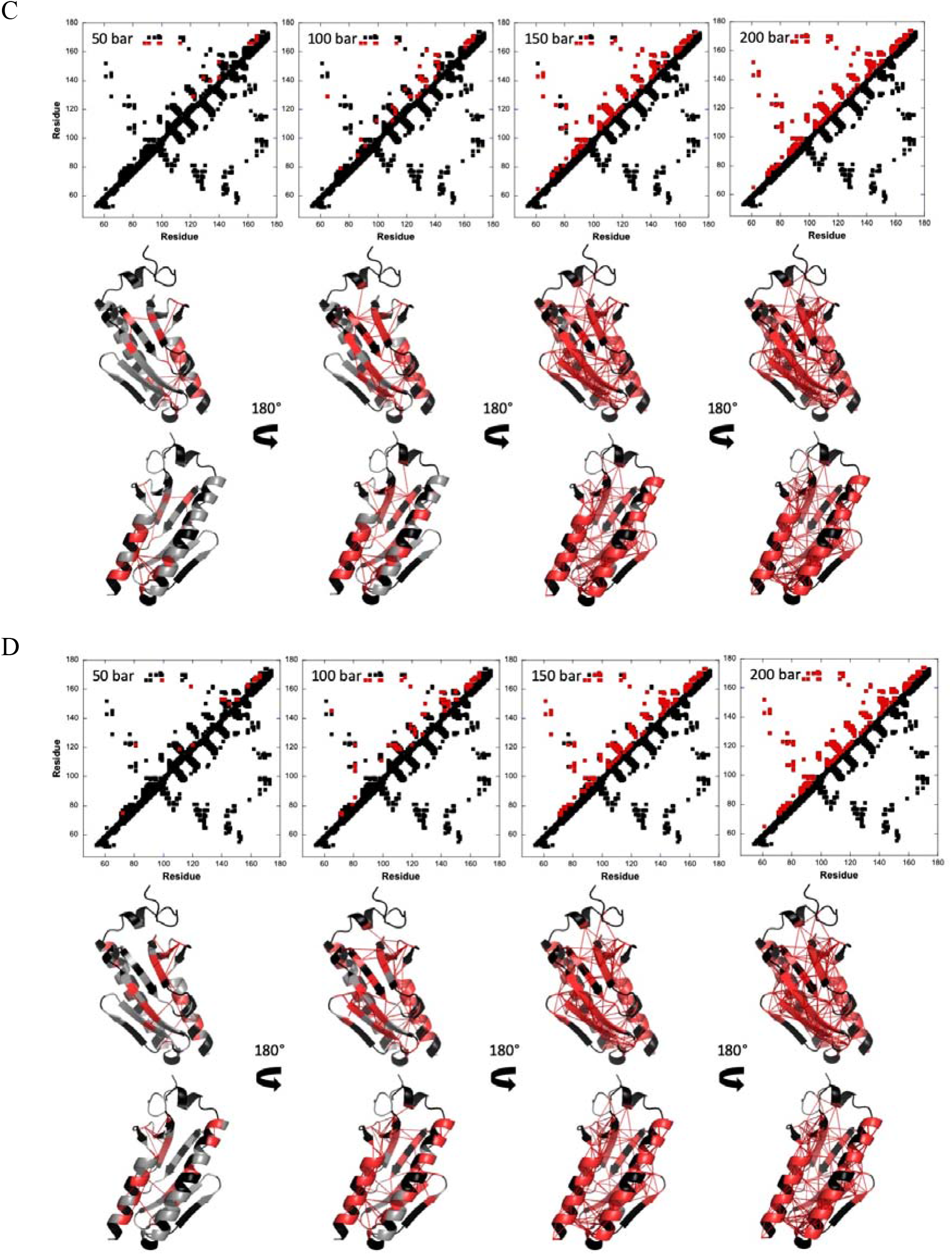
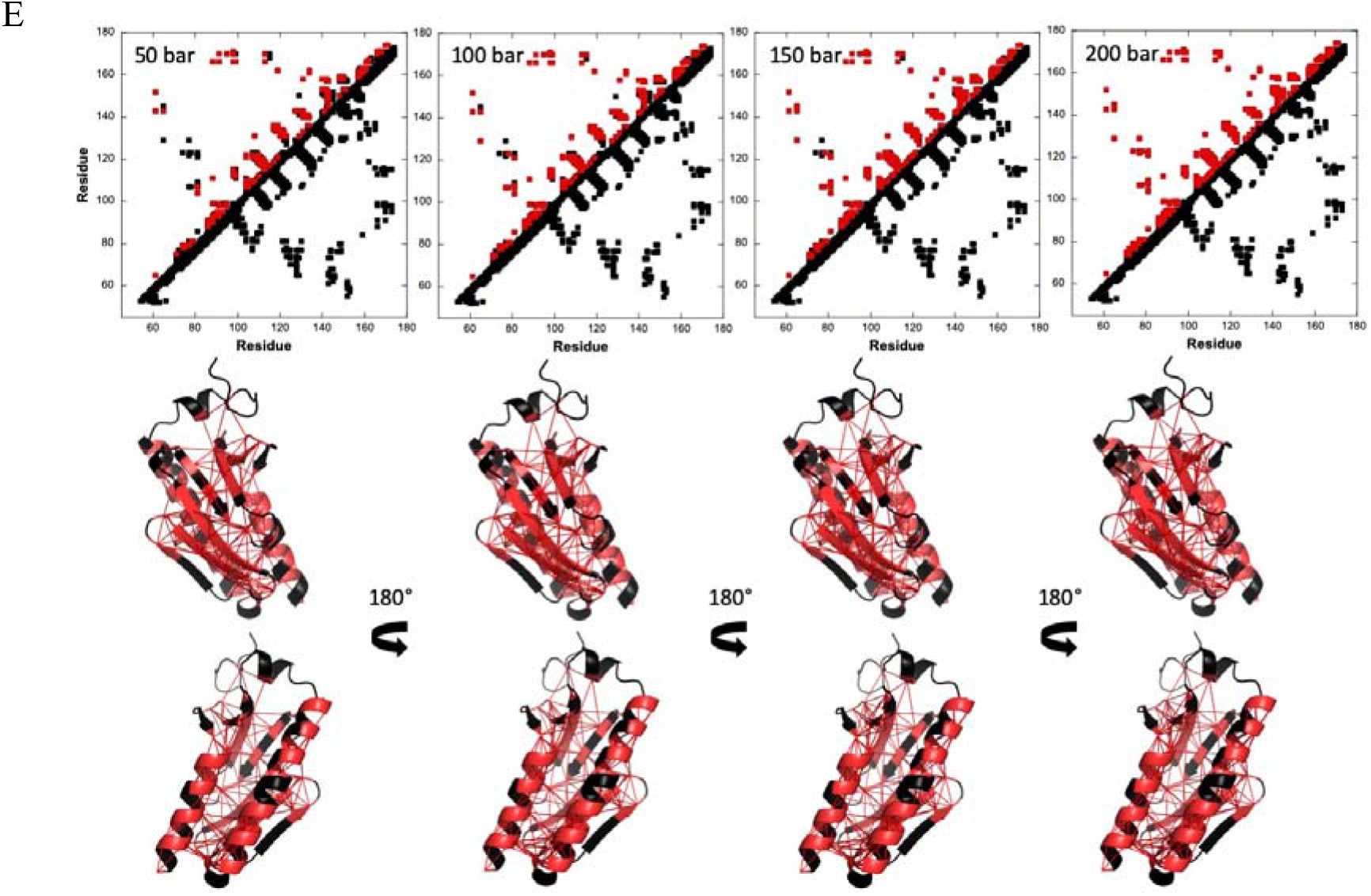
The pathway of pressure denaturation of Yfh1 at different temperatures and pressure. Data collected at (A) 283 K, (B) 288 K, (C) 293 K, (D) 298 K and (E) 303 K. Top) Fractional contact maps built from the AlphaFold model at 50, 100, 150 and 200 bar, as indicated. Contacts below the diagonal were calculated with CMview and correspond to residues where the distance between corresponding Cα atoms is lower than 9.5 Å. Above the diagonal, only the contacts (> i, i+1) for which fractional probability can be obtained have been reported. In addition, residues have been colored in red when contact probabilities *P*_*ij*_ lower than 0.5 are observed. Bottom) Visualization of the probabilities of contact on ribbon representations of Yfh1 (opposite views that differ for a 180° rotation along the vertical axis) at 50, 100, 150 and 200 bar. The red lines represent contacts that are significantly weakened (*P*_*ij*_ < 0.5) at the corresponding pressure. Residues involved in these contacts are colored in red, and residues for which fractional probability cannot be obtained are colored in black.

As a conclusion, these results indicate that temperature does not only modify the onset and the cooperativity of the unfolding transition but affects the unfolding pathway. This important observation suggests a different mechanism of unfolding for the high and low temperature transitions that is revealed at low pressure values, allowing us to follow the early stages of unfolding.

## Discussion

Here, we report a study of the properties under pressure-induced unfolding of Yfh1, a small globular protein from yeast and a member of the frataxin family highly conserved from bacteria to primates (Castro et al., 2019). Yfh1 is a unique model system that, thanks to its marginal stability, is particularly suited to study the mechanisms of protein unfolding under different denaturing agents (Pastore and Temussi, 2023). Pressure denaturation studies are interesting because they allow us to reach an important part of the phase diagram of protein unfolding that could not be accessed otherwise. They can also, in principle, access unfolding without the need of adding chaotropic agents such as urea or guanidinium that, inherently, perturb the chemical environment and changes the solvation state of proteins, acting as cosolvents (Pastore and Temussi, 2023). Finally, high-pressure studies provide a unique tool to assess cold denaturation, since high pressure increases the cold denaturation temperature and at the same time reduces the freezing point of water (Smeller, 2002).

In a previous study based on the intrinsic fluorescence of the two tryptophan residues present in the Yfh1 sequence, we had used pressure as a means of structural perturbation to obtain information on the global response of Yfh1 and describe the phase diagram of the protein (Puglisi et al., 2022). We had shown that Yfh1 is a system exquisitely sensitive to pressure since the protein can be pressure-unfolded at values appreciably lower than those required for unfolding most of the small globular proteins reported so far: it is sufficient to use pressures below 600 bar to achieve the practically complete unfolding of Yfh1 *versus* the often more than 2000 bar needed for other proteins (Roche and Royer, 2018; Smeller, 2002).

In the present study, we aimed at gaining information on the pressure unfolding pathways of Yfh1 at the residue-specific level by NMR: this technique is uniquely suited for protein unfolding studies at the residue-specific level, providing local information on the behaviour of different regions of a protein, as previously demonstrated in thermal unfolding studies (Puglisi et al., 2021; Puglisi et al., 2020). We recorded spectra in the range of 1-1000 bar and 278-303 K, although, technically, we could comfortably go below and above these pressures and temperatures (Kitahara et al., 2006). Previous studies had told us that most of the unfolding events for this marginally stable protein happen within these ranges of pressure and temperatures and we restricted our work to the range of temperatures at which this marginally unstable protein is not strongly destabilized at atmospheric pressure (Adinolfi et al., 2004), to be able to distinguish between the effects of pressure and temperature.

We observed a qualitatively similar pattern at all temperatures: according to our previous work (Puglisi et al., 2022), the unfolding pre-transition starts already at 50 bar. Around 400 bar, the molecule is almost completely unfolded with almost complete, but not total, disappearance of the well-dispersed resonances from the folded species. Some residual peaks from the folded structure remain also at 600 bar as it can be observed through the retention of resonances from the folded species in the spectrum, but the relative ratio accounts for less than 2% of the initial intensity, which is close to the noise level (data not shown). It is worth mentioning that we had observed the same minor retention of the folded spectrum also in NMR spectra recorded at low and high temperatures and at atmospheric pressure at values at which the CD spectrum of the unfolded species had reached a plateau (Pastore et al., 2007; Adrover et al., 2012). This indicates once again that NMR is a technique uniquely sensitive to detect even minute residual quantities of a species, under conditions when other techniques could not compete.

It is interesting to compare the thermodynamics parameters obtained from fluorescence and NMR measurements. The ΔG and ΔV values obtained in our prior work using high-pressure fluorescence (Puglisi et al., 2022) (**Table 1**) are significantly smaller than those reported here from high-pressure NMR. The discrepancy between both the ΔG and ΔV values could be explained by remembering that the intrinsic Trp fluorescence reports on variations of the local environment of the Trp side chains, which does not necessarily coincide with the global transition of the protein structure. Additionally, fluorescence spectroscopy observed all the states of the protein: unfolded, folded and potential intermediate states, while, in our exchange regime, NMR reflects at a residue level only the folded and unfolded states of Yfh1. Any multiple states populated during the unfolding, with possibly small ΔV, would be averaged in fluorescence resulting in ΔV values smaller than those measured by NMR. Accordingly, our fluorescence data seem to indicate at least a 3-states equilibrium (2 different slopes). It is also interesting to notice is that the thermal expansivity, that is the slope of the ΔV as a function of the temperature, is 2.30 ml/mol°C for Yfh1, that is, albeit larger, in the order of magnitude of values reported for proteins of similar size in other studies (1.15 ml/mol°C for GIPC10 (Dubois et al., 2021) and 1.71 ml/mol°C for pp32 (Fossat et al., 2016).

Overall, there are several important and novel conclusions of the present study that give us a new perspective of the effect of pressure on protein unfolding and the role that hydration plays in it. First, our data collectively hint at a higher similarity between the mechanism that governs cold and pressure denaturation over high temperature: the overall clear-cut low-field chemical shifting of the spectrum under these two conditions as compared to the high temperature unfolded spectrum clearly speaks in favour of a de-shielding of the amide protons, in agreement with a recent study on the effect of pressure on the chemical shifts of the cold shock protein B from *Bacillus subtilis* (*Bs*CspB) (Berner and Kovermann, 2024). Pressure-assisted cold denaturation using high-pressure quartz NMR tubes (Kitahara et al., 2006) had also hinted at a closer similarity of the pressure-assisted, and cold, and alcohol unfolded states, supporting the notion that, similarly to alcohol, also pressure and cold reduce the hydrophobic effect (Vajpai et al., 2013). This agrees with Privalov’s theory (1979, 1982, 1990) by which cold denaturation strongly depends on the higher affinity of water to apolar groups and on hydrogen bonding with the solvent, while heat denaturation is entropically driven resulting from increasing molecular motions. While our conclusions may have been suggested before, our work seems to be the first example in which experimental data directly indicate a similar role of hydration in cold and pressure unfolding.

Second, our analysis allowed us to follow the pressure, cold and heat unfolding pathways at the residue level and characterize the different unfolded states at different temperatures helped by pressure-assisted destabilization. Already in 1995, Jonas and cow. demonstrated the possibility to exploit pressure to assess cold denaturation of RNase A and compare cold, heat, and pressure unfolding states and showed a non-cooperative unfolding (Zhang et al., 1995). Later on, Babu et al. (2004) and Pometun et al. (2006) demonstrated the presence of a non-cooperative ensemble of conformations in the cold but not in the heat-denatured unfolded state of ubiquitin in reverse micelles at atmospheric pressure. This work consolidated the view that proteins, also under native conditions, exist not as a single conformation but as ensembles of interconverting, transient microstates. Following studies adopting an ensemble-based model of protein structure, for instance to characterize the denatured state of a whole database of human proteins (Wang et al., 2008), revealed important sequence-dependent thermodynamic properties of denatured ensembles as well as fundamental differences between the denatured and native ensembles (Hilser et al., 2006; Whitten et al., 2006). The possibility to follow ubiquitin under a variety of conditions confirmed that the pressure-assisted cold unfolding of ubiquitin is not a simple two-state process, and that several intermediates exist (Kitahara et al., 2006; Vajpai et al., 2013).

Our analyses allowed us to follow the hierarchical mechanism of Yfh1 unfolding. At 288 K, that is close to the maximal stability, pressure unfolding is highly cooperative, close to a global two-state equilibrium between the folded and unfolded populations of the protein. Out of this zone of stability, unfolding becomes less cooperative at low pressures, whereas at higher pressures, it is again highly cooperative at all temperatures indicating a sudden collapse of the structure and the opening of the hydrophobic core.

These observations should be put into the context of our previous work on the forces contributing to the stability of Yfh1. We have previously shown that the bacterial, yeast and human orthologs of Yfh1 share the same fold but have very different stabilities, with the yeast protein being marginally stable (Adinolfi et al., 2004). We proved a strong contribution of the C-terminus of the protein in stabilization: when we cut the much longer C-terminus of the bacterial or human proteins, we drastically reduced the Tm of these proteins, without affecting the low temperature transition (Adinolfi et al., 2004). When vice versa, we extended the C-terminus of yeast Yfh1 we gained in stability (ca. 10 °C in ΔTm). This is because the C-terminus inserts between the two helices and, without this insertion, the hydrophobic core is more easily accessible to the solvent. We then discovered that cold denaturation is detectable only for Yfh1 and only at low salt concentrations (Pastore et al., 2007; Sanfelice et al., 2014). We noticed that Yfh1 contains a superficial cluster of spatially close negatively charged residues (Sanfelice et al., 2015). We mutated up to three of these residues to neutral serines and found that these mutations lead to some stabilization at high temperature but to a much stronger effect at low temperature, making cold denaturation undetectable. We concluded that electrostatic repulsion is a strong element in inducing the cold transition of Yfh1 under conditions in which hydrophobic forces are weaker. In support to this hypothesis, we managed to convert the stable bacterial ortholog to a marginally stable protein that undergoes detectable cold denaturation by recreating the electrostatic cluster of the yeast protein through a few mutations (Bitonti et al., 2022). The data presented here perfectly agree with these previous conclusions as we observe that the pattern of unfolding upon increasing pressure depends on temperature: at moderate pressures, we observe two different temperature-dependent unfolding pathways: at low temperature (283 K), partial unfolding concerns primarily the N-terminal helix and the first strands of the ß-sheet, where the negatively charged cluster is (Bitonti et al., 2022; Sanfelice et al., 2015). At high temperature, unfolding affects first the C-terminal helix and then extends to the ß-sheet. This behaviour suggests a synergic mechanism between pressure- and temperature-induced denaturation in which pressure destabilization helps to unveil the hierarchical events of cold- and heat-induced unfolding.

Finally, it has been suggested that pressure denaturation depends on the elimination of the solvent-excluded internal voids and/or on the water penetration inside the core of the macromolecule (Roche et al., 2012). Accordingly, pressure denaturation studies provide an important thermodynamic parameter, that is the volume changes upon unfolding, ΔV. This parameter is defined as the difference between the system (i.e. the protein and the solvent) volumes of the unfolded and the folded states (or vice-versa reversing the sign). Over the temperature range of the present study, we consistently observed, as expected, that pressure denaturation shifts the equilibrium from the folded state to the unfolded state because the latter has a smaller system volume, leading to negative values of ΔV. The difference is persistently larger in absolute value at lower temperatures and becomes smaller at higher temperatures, as it was previously observed from high-pressure fluorescence studies on this protein (Puglisi et al., 2022), suggesting that increasing the temperature causes a more compact and less compressible state. The temperature-dependent decrease in the absolute value of 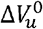 with increasing temperature is a well-known effect that has been ascribed to the difference in thermal expansion between the folded and unfolded states (Roche and Royer, 2018). However, we observe a consistent increase of the absolute value of 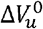 also at low temperatures at which the protein is becoming cold denatured (Puglisi et al., 2022; Mitra et al., 2008). This means that the thermal expansion of all protein states arises from the fact that hydration contributes to the system volume more significantly at lower temperatures due to stronger hydrogen-bonding, whereas waters will be increasingly more dynamic at higher temperatures, thus resulting in a lower contribution. It will remain to be seen whether the relationship is linear also at lower temperatures where the population of cold denatured increase.

In summary, our report describes the pressure unfolding of Yfh1 and allowed us to follow the unfolding pathway of the protein as a function of pressure and temperature in a residue-specific way. We could thus trace the regions of Yfh1 which unfold first and determine a model that could explain the mechanism of unfolding where we have observed an inherent interplay between pressure and temperature. The two main conclusions of our study are the higher similarity between the high pressure and the cold denaturation mechanisms, despite the inherently different physical processes, and, directly related to it, the determinant role of hydration in protein stability.

## Acknowledgements

We wish to thank Catherine Royer for helpful discussion and Stephen Martin for invaluable help with data interpretation.

## Notes

### Competing Interest Statement

The authors have declared no competing interest.

