## Supplementary material for "Understanding the physical determinants of pressure denaturation: Exploring the unfolding pathways of Yfh1 at different temperatures with High-Pressure NMR": Suppl. Figures S1 and S2

**Supplementary Materials**

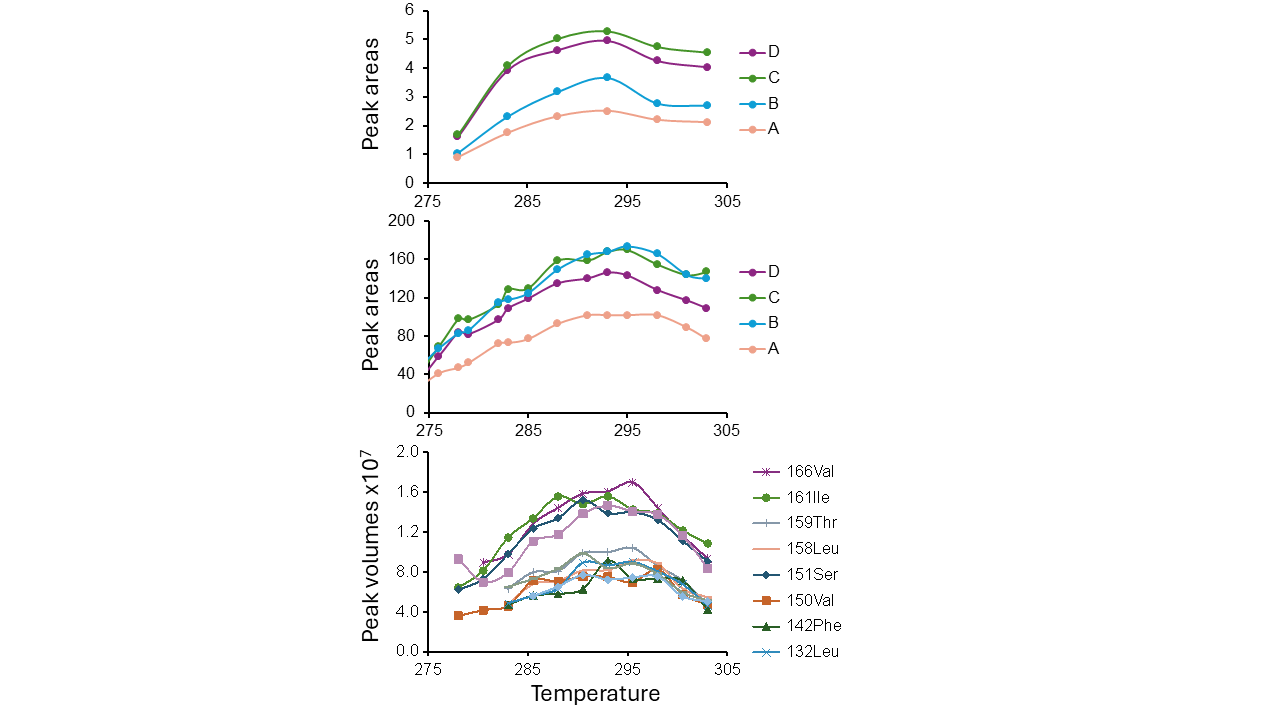

**Figure S1** – Dependence of the areas or volumes at room pressure as a function of temperature. Top: Plot of the areas of ring-current shifted resonances in the 1D spectrum as a function of temperature. Ring-current shifted resonances are particularly diagnostic of the protein core because they correspond to residues, typically methyl protons but not only, that are shifted as compared to their average position in a random coil protein because persistently sitting spatially close to aromatic groups (Perkins and Wuethrich, 1979). We had previously used these resonances to get the stability curves (i.e. the plot of ΔG versus temperature) at 1 bar, from which the thermodynamic parameters of the unfolding process can be obtained (Pastore et al., 2007; Martin et al., 2008; Puglisi et al., 2021). Using the assignment from BMRB (ID 19991), resonances correspond to A=ILE79 (HD1), B=ILE93 (HD2), C=ILE79 (HG13), and D=LEU85 (HD1). Middle: The same but from Pastore et al. (2007). Bottom: Peak volumes from 2D HSQC for the amide protons of specific residues that were identified as buried and thus being diagnostic of the unfolding process (Puglisi et al., 2019). The data were obviously collected on different protein preparations, produced by different researchers. The excellent agreement of the profiles gives us a measure of the reproducibility of the process.

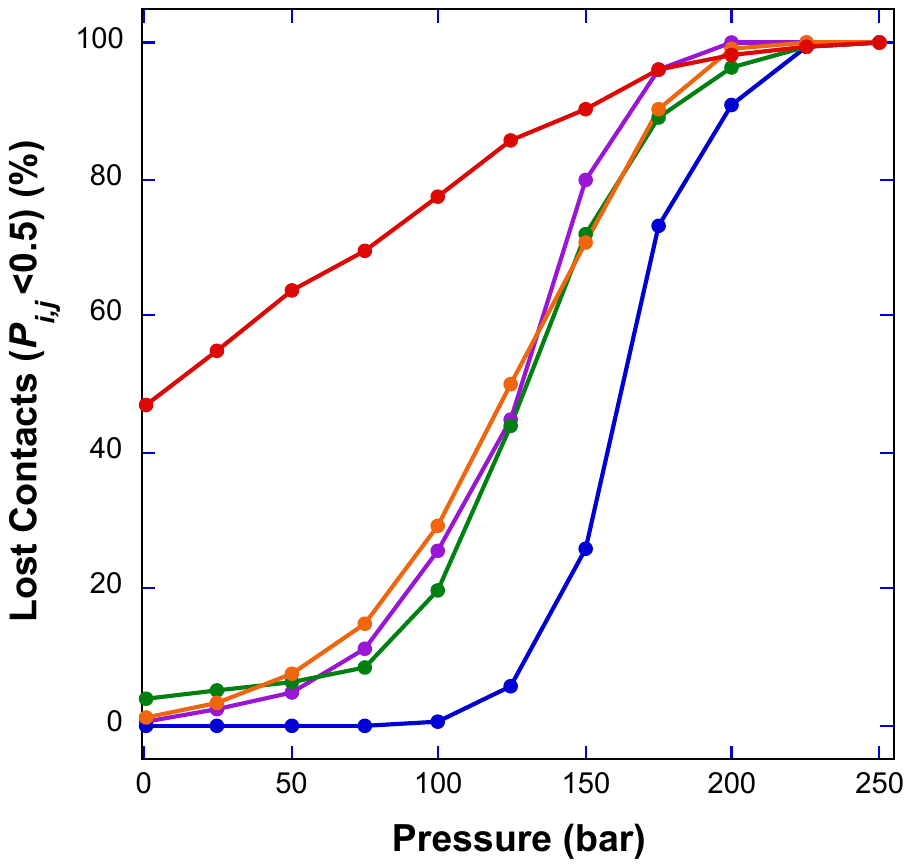

**Figure S2. Evolution of the Loss of Contact as a function of pressure during the unfolding transition of Yfh1.** The plot shows the data collected at 283 K (violet curve), 288 K (blue curve), 293 K (green curve), 298 K (orange curve) and 303 K (red curve). The considered contacts involve residues for which residue-specific pressure denaturation curves can be obtained at the five temperatures. A contact between two residues *i* and *j* was considered as lost when the corresponding calculated probability of contact *P_i,j_* was found lower than 0.5.
